# Cost-effective, high-throughput phenotyping system for 3D reconstruction of fruit form

**DOI:** 10.1101/2021.09.30.462608

**Authors:** Mitchell J. Feldmann, Amy Tabb

## Abstract

Reliable phenotyping methods that are simple to operate and inexpensive to deploy are critical for studying quantitative traits in plants. Traditional fruit shape phenotyping relies on human raters or 2D analyses to assess form, e.g., size and shape. Systems for 3D imaging using multi-view stereo have been implemented, but frequently rely on commercial software and/or specialized hardware, which can lead to limitations in accessibility and scalability. We present a complete system constructed of consumer-grade components for capturing, calibrating, and reconstructing the 3D form of small-to-moderate sized fruits and tubers. Data acquisition and image capture sessions are 9 seconds to capture 60 images. The initial prototype cost was $1600 USD. We measured accuracy by comparing reconstructed models of 3D printed ground truth objects to the original digital files of those same ground truth objects. The *R*^2^ between length of the primary, secondary, and tertiary axes, volume, and surface area of the ground-truth object and the reconstructed models was *>* 0.97 and root-mean square error (RMSE) was *<*3mm for objects without locally concave regions. Measurements from 1mm and 2mm resolution reconstructions were consistent (*R*^2^ *>* 0.99). Qualitative assessments were performed on 48 fruit and tubers, including 18 strawberries, 12 potatoes, 5 grapes, 7 peppers, and 4 Bosc and 2 red Anjou pears. Our proposed phenotyping system is fast, relatively low cost, and has demonstrated accuracy for certain shape classes, and could be used for the 3D analysis of fruit form.

## Introduction

Fruit appearance is a key trait for many crops and can condition market viability of fruit products and the success of cultivars (Costa et al., 2011; Gaston et al., 2020; Mathey et al., 2013). Taken together, the shape and color, or appearance, of fresh fruit are often associated with quality as they reveal condition and impact consumer perception of taste (Costa et al., 2011). Fruit shape is a heritable, complex trait that is difficult to assess due to the complex nature of data acquisition, which can be both time consuming and computationally laborious (Feldmann et al., 2020; Migicovsky et al., 2016a; Turner et al., 2018; Visa et al., 2014; He et al., 2017; Minervini et al., 2015; Atefi et al., 2021; White et al., 2000). Fruit shape in agricultural studies have primarily been assessed subjectively, placing fruit into qualitative bins ranging from ‘deformed’ to ‘uniform’ or by using 2D geometric morphometrics (Ishikawa et al., 2018; Visa et al., 2014; Feldmann et al., 2020; Migicovsky et al., 2016b; Horgan, 2001; Horgan et al., 2001; Wu et al., 2015, 2018; Monforte et al., 2013; Liao et al., 2018).

Publicly available methods for 2D phenotyping of plants and plant organs have increased in recent decades to support high quality analysis of leaves, roots, shoots, stems, tubers, and fruits (Seethepalli et al., 2018, 2020; Topp et al., 2013; Balduzzi et al., 2017; Turner et al., 2018; Salungyu et al., 2020; Li et al., 2018a,b; Warman et al., 2021). Computer vision has shown great potential to quantify external fruit quality and 2D imaging has been successfully implemented to measure the shape and size of fruits such as strawberries (Ishikawa et al., 2018; Feldmann et al., 2020), apples (Migicovsky et al., 2016a), carrot (Horgan, 2001; Turner et al., 2018), mangoes (Naik et al., 2015), and many others. More recently, methodologies for 3D reconstruction of plant organs have been developed with approaches that vary in speed, scale, cost, and accuracy; including laser scanners, x-ray computed tomography, and reconstruction from sequences of 2D images from digital cameras (Gaillard et al., 2020; Chaudhury et al., 2020; Feldman et al., 2021; Dutagaci et al., 2020; Artzet et al., 2020; Teramoto et al., 2020; He et al., 2017; Paulus et al., 2014; Li et al., 2014; Liu et al., 2020; Dowd et al., 2021; Jiang et al., 2019; Sandhu et al., 2019; Hu et al., 2020). Methods that rely on sequences of 2D images are numerous and variable with their own complexities and nuances that provide different strengths and weaknesses (He et al., 2017; Porter et al., 2016; Sandhu et al., 2019; Hu et al., 2020; Wang et al., 2019; Liu et al., 2017, 2020; Warman et al., 2021).

Modern technologies and analyses can be used to assess these physical characteristics and ultimately provide researchers with the tools necessary to support genetic inquiries and biological discoveries, expand what is known about modern germplasm, and, potentially, enhance breeding practices for fruit form in fruit and vegetable crops (Klingenberg and McIntyre, 1998; Bookstein, 1997; Kuhl and Giardina, 1982; Chitwood, 2014; Chitwood and Otoni, 2017; He et al., 2017; Li et al., 2020; Feldmann et al., 2020; Turner et al., 2018; Gage et al., 2019; Ubbens et al., 2019). Multivariate and spatial statistics can be used to determine parameters that identify and quantify fruit defects (Wang et al., 2021), differentiate between marketable and non-marketable fruit (Ishikawa et al., 2018; Li et al., 2020), and understand fruit phenotypes that impact markets requiring long shelf-life and sustained fruit quality through harvesting, handling, and shipping.

This paper describes a rapid (9 s), low-cost ($1,600), turntable-type system for 3D reconstruction of fruit and tubers. Fruit rotates on an automated pedestal while a remote-controlled digital camera acquires images, as shown in Figure 1. We use a multi-camera calibration method (Tabb et al., 2019) to compute the calibration parameters of the camera at every time step. Fruit are segmented from non-fruit regions in the images. Finally, a reconstruction method using silhouettes as features (Tabb, 2013) reconstructs the fruit or vegetable shape using the calibration and silhouette information (Figure 2). A preliminary version of this work appeared as an extended abstract (Feldmann et al., 2019).

**Fig 1.**
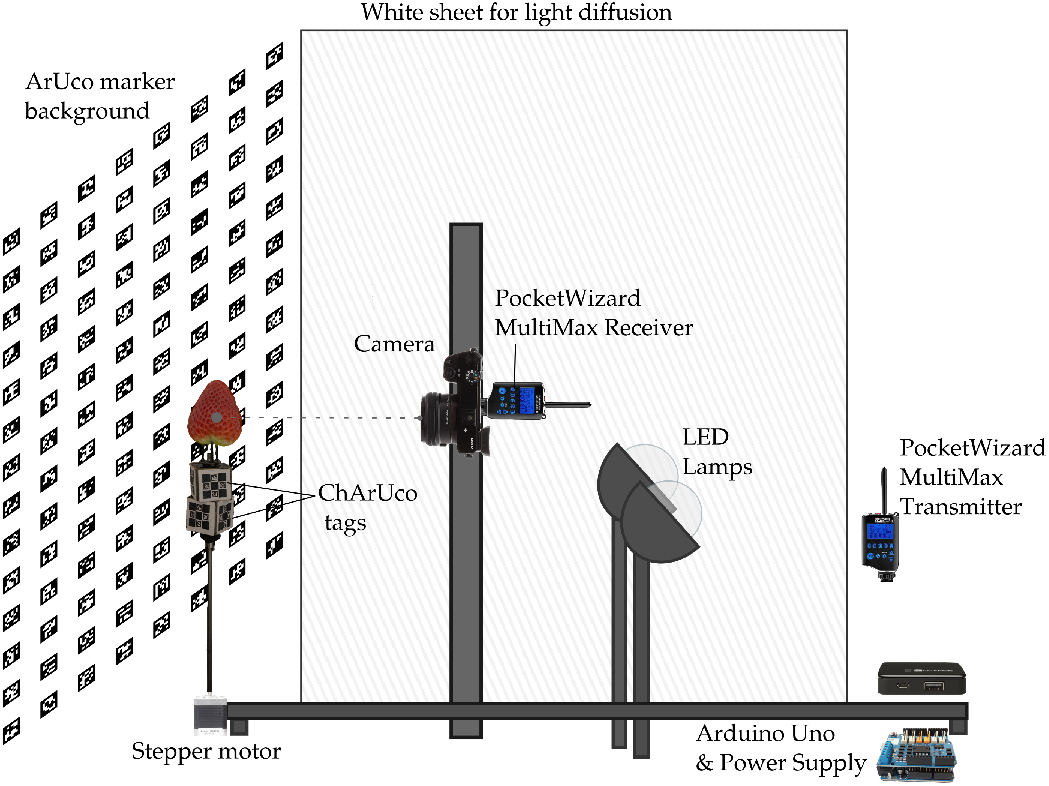
Imaging system hardware. From left to right: the arUco marker backdrop; a stepper motor with a metal pedestal, chArUco tagged cubes, and a target object, a strawberry; the 80/20 t-slotted aluminum inverted T () frame; a digital camera is mounted on the vertical limb of the frame and attached to a PocketWizard MultiMax II radio transceiver; reverse facing LED light sources; Arduino microcontroller connected to stepper motor and power supply. **Best viewed in color**.

**Fig 2.**
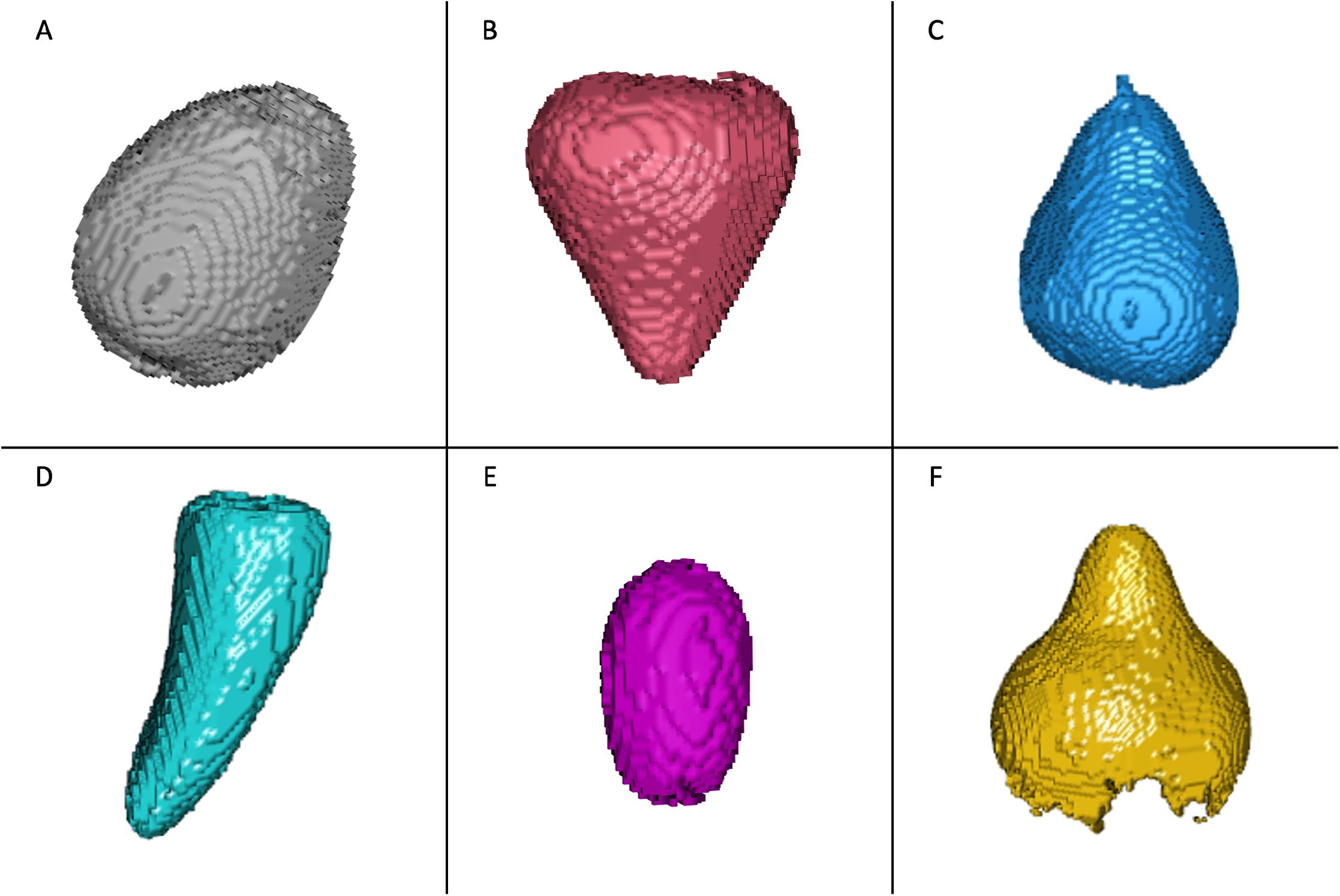
Representative model reconstructions of the six types of fruit or tubers imaged in this study. The resulting 3D reconstructed models from the hardware and software presented in this study, including: (A) a baby yellow potato, (B) a strawberry, (C) a Bosc pear, (D) a sweet mini pepper, (E) a green table grape, and (F) an Anjou pear where the segmentation failed All models shown are at 1mm resolution. Images are not to scale. **Best viewed in color**.

### Contributions

Our contributions to the state-of-the-art in fruit phenotyping and estimating 3D reconstructions are a high-throughput (9 second data acquisition), modular reconstruction system that can be used in lab or field settings (on a table) with high accuracy for objects that do not have local concavities. Our work is most similar to that of Sandhu et al. (2019) and He et al. (2017). In Sandhu et al. (2019), a turntable system is used and the cameras rotate around the target object, rice inflorescences. Relative camera calibration parameters are estimated by detecting features and estimating matches from color checkerboard pages and a structure from motion approach generates point clouds. This approach works well for the target crop, but has an unknown scale factor that is not solved for. Consequently, the physical units such as a mm or cm are unknown, and comparison with another system of a different size may be difficult. In our system, we calibrate directly from patterns in the scene, so the physical size of the sample is estimated. He et al. (2017) uses a turntable system in the configuration that we do, where the fruit is rotating and the camera is stationary, to acquire images for 3D reconstruction. They use a commercial software package to reconstruct point clouds, and then process point clouds to extract fruit features. Our work differs from both of these in that we use silhouettes and directly use voxel representations instead of point clouds.

### Core Ideas

- A low-cost 3D fruit phenotyping system is presented.
- Image capture using the proposed approach lasts for only 9 seconds.
- Accuracy is measured against 3D printed ground-truth objects.
- Camera calibration, background segmentation, and reconstruction does not rely on commercial software.
- An RMSE less than 3mm was obtained for ground truth objects without locally concave regions.

## Materials and Methods

The 3D phenotyping system consists of multiple parts: the physical system for data acquisition and the algorithms for reconstructing shape from that data. Briefly, one or multiple digital cameras are mounted on an aluminum frame and remotely triggered at a frame rate of 7 frames per second (FPS) for 9 seconds while a stepper motor controlled by a microcontroller rotates an object. We do typically only refer to the session as the time elapsed between triggering cameras and stepper motor and the time that the cameras stop capturing, as the time between objects is variable depending on hardware availability and user experience. Captured images are calibrated using CALICO, a multi-camera calibration method (Tabb et al., 2019) that relies on a combination of arUco and chArUco markers (An et al., 2018; Garrido-Jurado et al., 2016; Romero-Ramirez et al., 2018). The fruit foreground is segmented from the non-fruit background in each calibrated image. Segmented silhouettes of fruit are then used as features to reconstruct 3D models.

### Hardware

The physical system is composed of an aluminium frame, digital cameras and camera control units, a USB barcode scanner, a microcontroller, and a microcomputer.

#### Frame

The frame’s design is an inverted “T” shape structure with a 1.22m (4ft) horizontal base and a 1.22m vertical arm (Figure 1). The main structure is composed entirely of 80/20 t-slotted aluminum. We chose this material because it is lightweight, strong, and inexpensive. The t-slot design and the availability of different fasteners provides rigidity while remaining modular. The arm is connected to the base using a side-mount and hand brake. The side-mount and hand-brake combination means that the vertical arm can be positioned anywhere along the length of the horizontal base. The camera is mounted to the vertical arm using the same side-mount and hand brakes, again allowing it to be positioned continuously along the vertical span of the arm.

#### Cameras and controllers

We used one Sony *α*6000 mirrorless digital camera for this project. The camera was set to medium speed continuous image capture (7 frames per second), manual focus, and aperture priority mode with the aperture set to f/8. We controlled the camera with a PocketWizard MultiMax II transceiver unit. These units attach to the camera’s multi-port and digitally control the camera’s shutter button and can be programmed to “hold” the shutter button to allow for variable duration. We used 9 seconds of hold to match the rotation rate of our stepper motor. Two PocketWizard MultiMax II units are required: one unit to transmit a signal and one unit to receive a control signal for multiple cameras. With these, the camera is controlled from a single source which is triggered by the input of a barcode scanner.

#### Microcomputers and stepper motor

The data acquisition process consists of rotating the fruit on the pedestal and acquiring images of that fruit. To automate this process, we used a Raspberry Pi 3 microcomputer as well as an Arduino Uno Rev3 microcontroller. To rotate the objects on the pedestal, we used a Nema 17 stepper motor controlled using an Arduino Uno Rev3 and an Arduino Rev3 motor shield. The pedestal is a thin metal rod approximately 20cm in length and 5mm in diameter. The Nema 17 stepper motor has 200 steps per rotation (1.8° per step). The motor is programmed to take 1 step every 45ms, which is a full rotation every 9 seconds.

#### Lighting

We used 4 LED lamps to illuminate the scene. These lights are all directed away from the object towards a reflective white sheet to reduce the intensity of the light on the scene. This enabled us to dramatically reduce, and in some instances eliminate, the glare on the surface of more reflective objects such as strawberries. The lights chosen do not have any temperature control and are likely not ideal for color accurate measurements.

#### Calibration Targets

Calibration is performed on image data that also contains the data for reconstruction. To accomplish one-step calibration and data acquisition, the workspace is prepared with calibration targets, which are shown in Figure 1. The fruit or tuber is mounted on the pedestal. A pair of offset cubes are mounted on the pedestal directly below the fruit or tuber (An et al., 2018). Each cube is *×* 2.5cm *×* 2.5cm *×* 2.5cm with a small hole in the center for mounting onto the pedestal, and are rigidly attached.

On the cube faces without holes, chArUco markers are printed and attached to the cubes. The chArUco patterns are a 3 *×* 3 checkerboard with each square unit measuring 6.67mm×6.67mm. The two cubes have eight faces with chArUco patterns on them, and multiple cube faces should be visible in any frame providing enough information such that the calibration method CALICO can compute camera poses. We use a scene background, a 0.71m^2^ (26in^2^) aluminum panel, composed of arUco markers (Garrido-Jurado et al., 2016; Romero-Ramirez et al., 2018). This type of background allows us to refine internal camera calibration parameters using the multi-camera calibration method, CALICO. Each arUco marker is 2.25cm^2^ and adjacent markers are separated by 2.75cm of white space. Each image contains between 60-70 unobscured arUco markers, depending on the size of the object.

#### Data Acquisition control

For every object, the Arduino board is first activated by the user, who starts the stepper motor’s rotations. The cameras are triggered once the motor begins to rotate using a barcode scanner and a custom python script that sends a serial signal through the general-purpose input output (GPIO) of the Raspberry Pi to the transmitter unit. The cameras stop firing after 9 s and Arduino board is deactivated manually. Including swapping objects, the time for each sample is less than 25 s. In this time period, each camera captures approximately 60 frames during one complete rotation of the object. Following each session, the camera must be allowed to clear its on-board cache and to write the images to an SD card. Depending on the camera and the SD card, write speeds may vary. In our setup, this typically took about 10 s.

### Reconstruction Pipeline

The 3D reconstruction pipeline consists of three stages: calibration, segmentation, and reconstruction. In this work, the stages consist of independent modules but they have been selected based on the assumptions of the reconstruction module.

We use a Shape from inconsistent Silhouette method from Tabb (2013) (Tabb, 2013) that requires camera calibration information and uses silhouettes – or segmentations – of the target object to generate reconstructions. This reconstruction method tolerates calibrations with small camera calibration error and small image segmentation error. The methods used for calibration and segmentation are discussed in the Calibration and Segmentation sections, respectively.

#### Calibration

The reconstruction method depends on camera calibration. Camera calibration usually means the knowledge of the internal camera parameters as well as the external pose (rotation and translation) of cameras relative to a world coordinate system. In the context of this work, by ‘computing the calibration,’ we mean determining the internal camera parameters as well as the external pose of the camera with respect to the calibration object at each image acquisition.

We use an existing method for multiple-camera calibration, CALICO (Tabb et al., 2019), to compute the desired calibration parameters. To use CALICO in this context, chArUco tags have to be rigidly attached to the pedestal and multiple tags visible at each time instant, which is why the physical system is prepared as in the Hardware section. Some datasets had significant error in the camera pose because not enough chArUco tags were detected in each frame, so we extended the ‘rotation’ option of CALICO to also detect and use the arUco corners within the chArUco boards.

A successful calibration estimates internal camera calibration parameters, selects one of the chArUco boards as a world coordinate system, and estimates the relative pose of the camera at each image acquisition with respect to that world coordinate system. For instance, see Figure 4, which shows an example of an input image, (Figure 4(A)) and the chArUco markers below the pedestal. The reconstructed chArUco board poses, camera poses at each image acquisition, and a reconstructed strawberry are shown in Figure 4(C).

**Fig 3.**
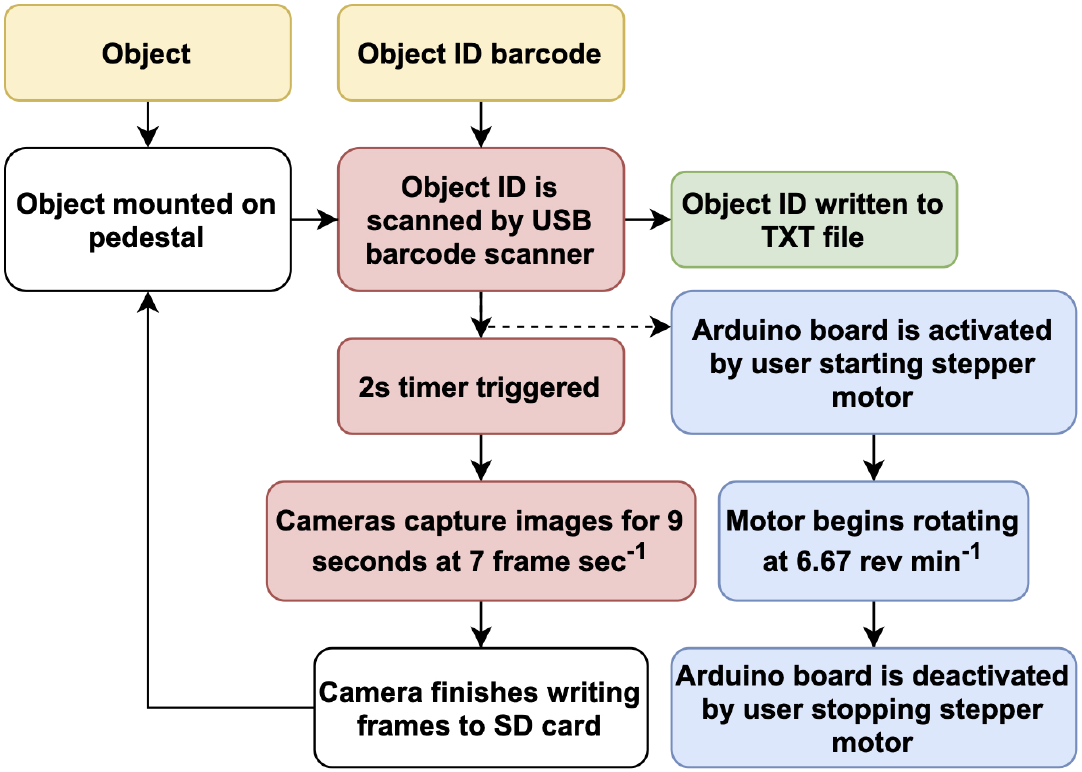
Flow diagram of data acquisition strategy and steps. Input (yellow): A physical object with an associated barcode ID, e.g., QR code or data matrix. Staging (white): The object is placed on the pedestal. Camera Triggering (red): The object ID barcode is scanned using a USB barcode scanner attached to a Raspberri Pi, starting a 2 second timer before triggering the cameras for 9 s at 7 FPS. Intermediate output (green): The scanned barcode is written into a TXT file. Motor control (blue): During the 2 second timer, the Arduino board is activated by supplying power to the board initiating rotation of the object at 6.67 RPM (1 revolution every 9 seconds). Once the image capture is complete, the user deactivate the Arduino board, stopping the motor. Staging (white): The camera will need upwards of 15 seconds to finish writing the images to storage, during which time the next object can be staged on the pedestal, initiating the following session. During this process, the user is responsible for mounting the objects to the pedestal, scanning the barcode ID, and activating/deactivating the stepper motor. **Best viewed in color**.

**Fig 4.**
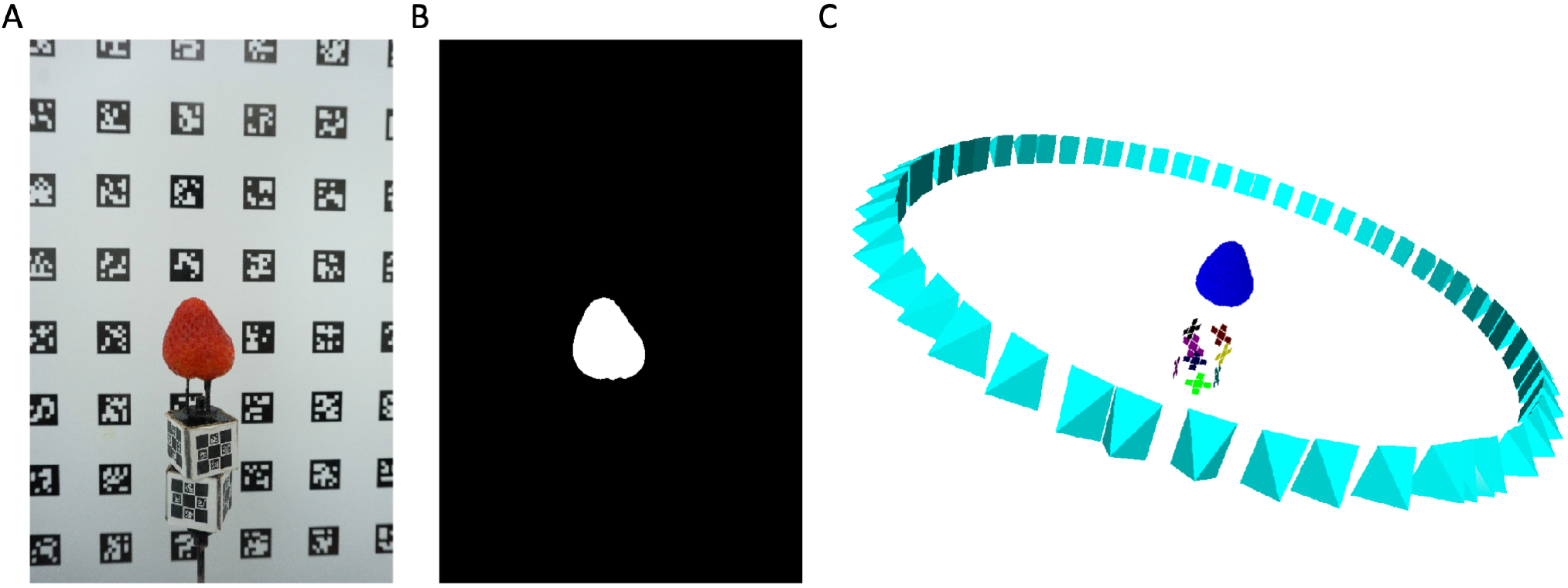
An example of successful camera calibration results. Overview of image data and results. Images of fruit and calibration objects are captured while the pedestal rotates (A). Each image is segmented (B). The calibration and segmentation information is used to reconstruct the fruit shape, 1 mm voxel resolution shown in (C). A camera pose is represented as a pyramid, where the camera center is the tip of pyramid. Three-dimensional visualizations for this figure, Figure 5, and Figure 7 generated with Meshlab (Cignoni et al., 2008). **Best viewed in color**.

**Fig 5.**
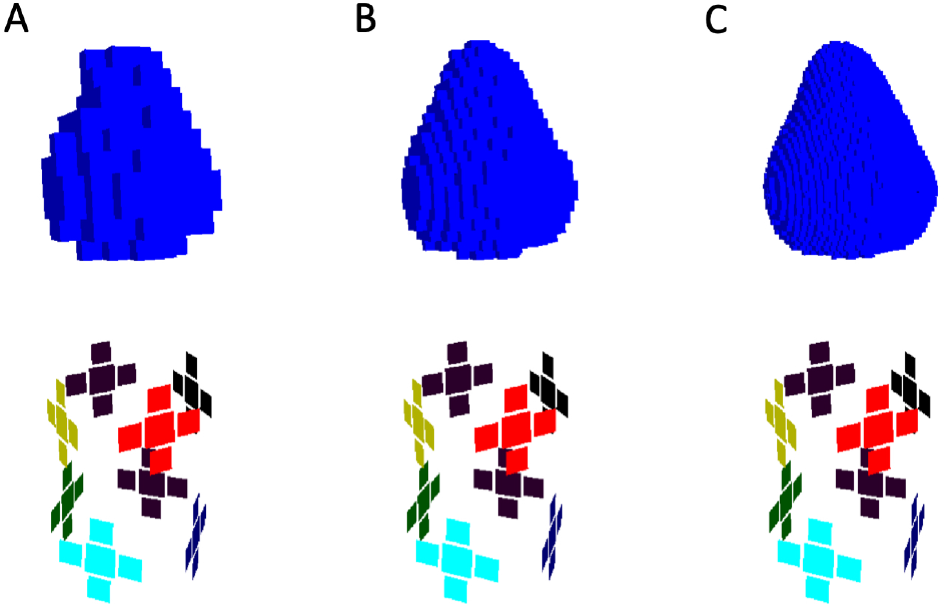
Reconstructions of a strawberry at 3 different resolutions. Strawberry reconstruction from Figure 4 during the hierarchical reconstruction process, with estimated calibration pattern positions below. The SfIS reconstruction method starts at a large voxel resolution (here, 4 mm) (A), and refines the reconstruction at finer resolutions using the prior level’s results (B) 2 mm, and (C) 1 mm. **Best viewed in color**.

#### Segmentation

The reconstruction method depends on segmented images, where the fruit, tuber, or ground truth object is separated from the image background as in Figures 4 and 6. The backgrounds consist of arUco tags and the pedestal with chArUco tags. With this background, consisting of dark and light intensities, we took an approach of modelling the actual intensities of the calibration tags per image as a Gaussian Mixture Model with two components, and then used background subtraction to determine the location of target objects. First, the arUco tags are located in the image. Then, the dark and light regions of each tag are identified using Otsu’s segmentation algorithm (Otsu, 1979). The dark and light regions of all of the tags are used to estimate six Gaussian distributions (masks are shown in Figure 6): 𝒩_*d,r*_(*μ, σ*^2^), 𝒩_*d,g*_(*μ, σ*^2^), 𝒩_*d,b*_(*μ, σ*^2^), the distributions representing the dark intensities for red, green, and blue channels, and the same for all three channels of the light intensities.

**Fig 6.**
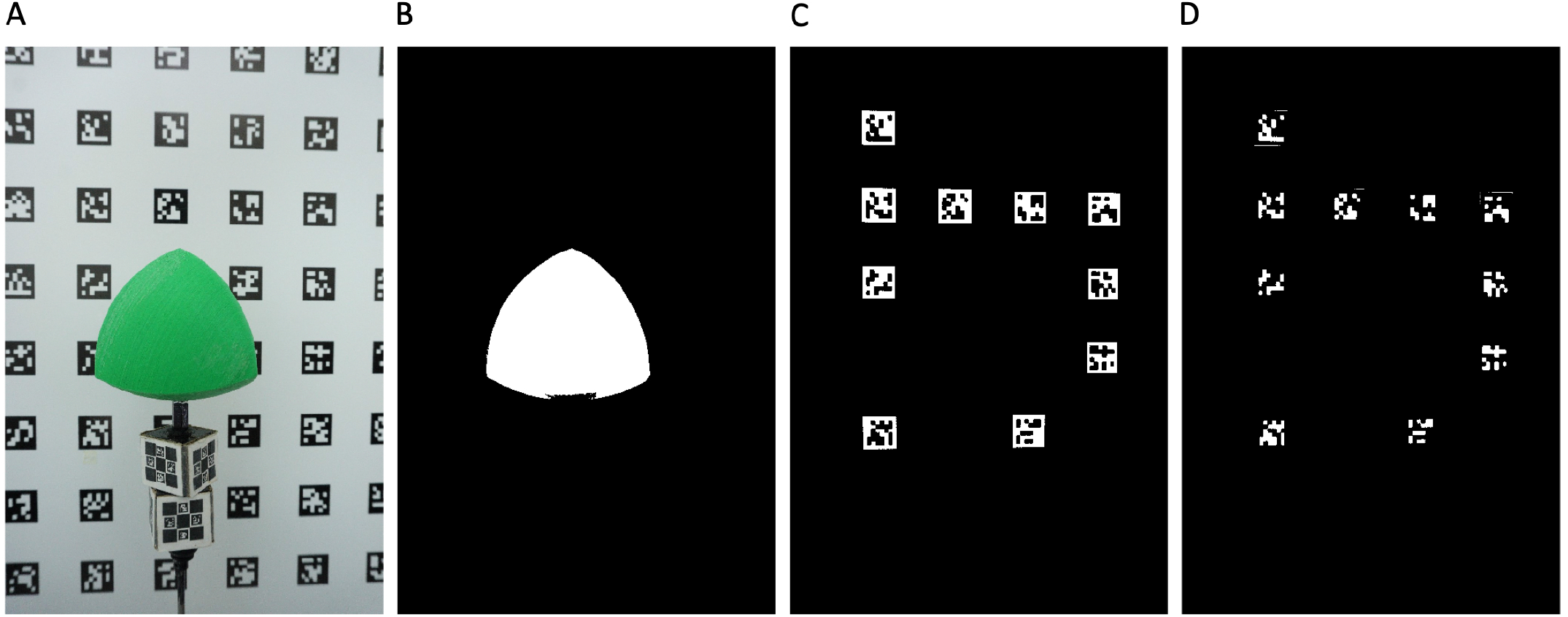
An example of successful background segmentation. Segmentation process from sample 1, one of the 3D printed objects. arUco tags are detected in individual images in the data acquisition step (A). Background subtraction generates the segmented image (B). Individual tags are segmented to separate the dark (C) and light (D) intensities; these regions are used to model the background. Notice a small segmentation error from shadow on the bottom of the tetrahedron. See text for more details. **Best viewed in color**.

Each image pixel *x* is evaluated against the distributions as in Equation 3. We use a typical background subtraction technique in that we subtract the mean and compare with a threshold; here the threshold is a constant multiplied by the standard deviation (see Piccardi (2004) for a review of background subtraction). The user provides constants *k*_*d*_ and *k*_*l*_, and from the distributions computes Boolean values *y*_*d*_ and *y*_*l*_ for each pixel *x*. The segmentation result of whether the pixel represents the background (0) or not (1) is stored in *z* = *y*_*l*_ *∧ y*_*d*_.

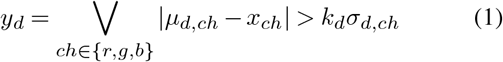

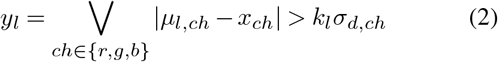

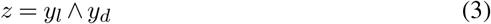

mIn our experiments, *k*_*d*_ = 2.0 and *k*_*l*_= 2.5 for all tests.

#### Reconstruction

We used a Shape from Inconsistent Silhouette (SfIS) method (Tabb, 2013) for 3D reconstruction of the plant organs and ground truth objects. With camera calibration and segmentation or silhouette provided, SfIS is a voxel-based method that searches for a labeling of voxels as occupied or empty such that the voxels match the provided segmentations. The match does not need to be exact, so some small camera calibration and segmentation errors can be present.

A key feature of the SfIS method is that it will not reconstruct concavities in 3D space. As examples of these types of shapes, the tetrahedron, sphere, and F ground truth objects (Figure 7) can all be reconstructed because they do not contain concavities, while the 6-sided spherical die cannot (Figure 7). The stem or calyx region of an apple is also an example of a locally concave region on a surface. The reason that the SfIS method is not able to reconstruct locally concave regions if because of its dependence on segmentations as features.

**Fig 7.**
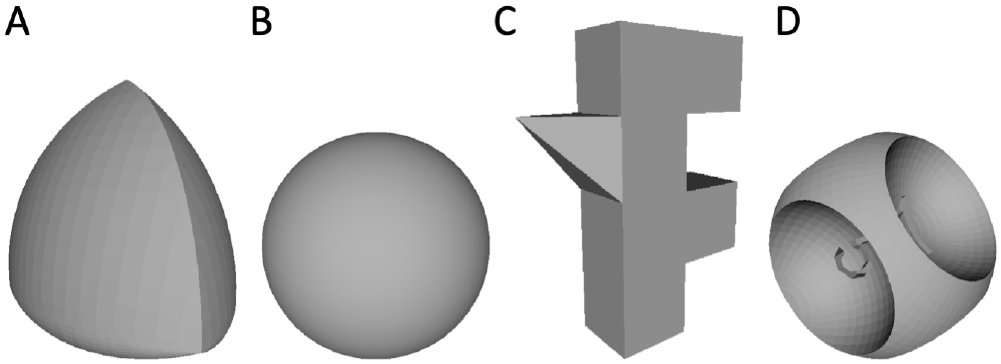
Ground truth objects. The four classes of ground truth objects used in this study. The 3D model files were used to print out physical copies, which were then imaged with our phenotyping system and reconstructed. (A), (B), and (C) can all be reconstructed with our system because they do not have locally concave regions, while the depressions in (D) cannot.

We use the extension to SfIS of hierarchical search described in Tabb (2014); the user specifies an initial voxel size, finds a solution with SfIS, divides the voxel size by eight and continues with search with SfIS, using the previous voxel size’s result as an initial solution. In this work, we performed experiments on all of the samples with two different parameter sets. In the first, the initial voxel size is 4 mm, the number of voxel divisions is two, and the final voxel size i s 2 mm. In the second set of experiments, the initial voxel size is 2 mm, the number of voxel divisions is two and the final voxel resolution is 1 mm. An additional parameter is the factor that the the input image is resized down, that value is 4 for both experiments. The initial image size is 6000 by 4000 pixels.

### Experiments

We focus on two primary experiments. The first is to quantitatively measure and evaluate our 3D model reconstructions against objects with a known shape. These objects with a known shape are the ground truth samples, with 3D model files that are 3D printed. The second experiment is to qualitatively assess the system’s ability to reconstruct various fruit models across different scales. We reconstructed the objects in both experiments at 1mm and 2mm resolution.

### Ground Truth Samples

Fruit form, especially in 3 dimensions, can be difficult to quantify. To assess the accuracy of our system, we selected shapes for which we had 3D model files, printed those files, and then reconstructed the models from image data with the phenotyping system. Through this process we can characterize the performance of our method on reconstructing different shape types with durable objects versus individual fruit measurements, where the fruit decays quickly and the human-made measurement cannot be precisely replicated.

A motivation for using 3D printed objects is to have a way to quantitatively assess the performance of the phenotyping system, with a durable artifact that can be stored indefinitely and re-printed and/or scaled if needed. Since we have the originating 3D model file, we can compare the reconstruction and the ground truth object in ways that human-made measurements are unable to, by assessing differences in surface area and volume. This is in contrast to measurements on fruits or tubers that will not persist past a single session and may suffer from measurement error.

We identified 4 digital objects from Thingiverse (https://www.thingiverse.com) that had good representation of many different shapes that are both common and uncommon in 3D biological structures, such as fruit and tubers: convex regions, saddle regions, and locally concave regions, shown in Figure 7. We scaled these 4 objects prior to printing so that we would have different size representations. We 3D printed these 11 object *×* scale stereolithography (STL) format files (Feldmann and Tabb, 2021e,d,c,b) using a commercial-grade 3D printer. The 3D printed objects were then imaged in our system and reconstructed from the 2D images.

### Quantitative Analysis

Once the ground truth objects are reconstructed, some postprocessing is done to align the reconstructed 3D model with the digital ground truth 3D model files. Specifically, we used R 4.1.0 (R Core Team, 2019) to perform quantitative comparisons between ground-truth objects and reconstructed models with the packages Morpho and Rvcg (Schlager, 2017), rgl (Murdoch and Adler, 2021), Lithics3D (Pop, 2019), and mesheR (Schlager, 2020). The reconstructed models are in Polygon (PLY) file format and were read using *Rvcg::vcgPlyRead()*. STL objects, e.g., the ground-truth objects, were read using *rgl::readSTL()* and converted to mesh3d using *rgl::as*.*mesh3d()*. The *mesheR::icp()* function was used to perform the iterative closest point algorithm between the reconstruction and the ground-truth triangular meshes, with 100 iterations and allowing for reflection.

For these quantitative analyses, we chose to measure the magnitude of the primary, secondary, and tertiary axes, e.g., X, Y, and Z, the surface area *SA*, and the volume *V ol*. the difference in magnitude between two models, *δ*_*X*_, *δ*_*Y*_, and *δ*_*Z*_, are calculated as the difference between the magnitude of the first, second, and third axes of the reconstruction and ground-truth following ICP alignment:

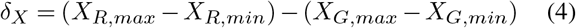

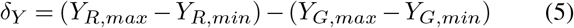

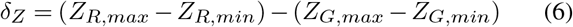

where *X*_*G,min*_ and *X*_*G,max*_ are the minimum and maximum value of the first dimension of the ground-truth object *G*, respectively, and *X*_*R,min*_ and *X*_*R,max*_ are the minimum and maximum value of the first axis of the reconstructed object *R*, respectively. The *Morpho::meshDist()* function was used to calculate and visualise distances between 3D objects. The distance of the reconstructed model from the ground truth is summarized using root mean square error (RMSE). RMSE is calculated as:

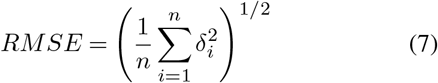

where *δ*_*i*_ is the distance between *i*-th pair of *n* corresponding points on the surface of the reconstruction and ground-truth objects. The volume and surface area of models was extracted using *Lithics3D::mesh_volume()* and *Lithics3D::mesh_area()*, respectively The *rgl::shade3d()* function was used to visually compare 3D objects. All regressions were performed using *stats::lm()*.

### Sample Collection

We purchased fresh fruit and produce from a local grocery store in Davis, CA, USA for qualitative assessment. In total we purchased, scanned, and reconstructed 48 objects; including 18 strawberries, 12 potatoes, 5 grapes, 7 peppers, and 4 Bosc and 2 red Anjuo pears. We want to test our approach for robustness, and so chose fruit and produce with different scales, colors, levels of glossiness, and other features.

### Qualitative Comparisons

Reconstructed fruit were oriented using principal components analysis (PCA) with the *stats::prcomp()* function. PCA orientation of the 3D reconstructed models orients results in 3 axes corresponding to the primary (X), secondary (Y), and tertiary (Z) axes ordered by magnitude, e.g., X ≥ Y ≥ Z. PCA oriented models were visually inspected using *rgl::shade3d()* functions. Quantitative measurements (Volume, Surface Area, X, Y, and Z) were measured from the PCA oriented reconstructed models.

## Results

### Overall assessment of platform

Our combination of hardware and software was able to accurately reconstruct models of fruit (Figure 2) and ground-truth objects (Figures 8 and 9; Table 1). Image acquisition occurs in 9 s sessions and is buffered by approximately 15 seconds while the cameras finish writing photos to storage and the following object is staged on the pedestal (Figures 1 and 3). It is possible to achieve about two sessions per minute with current parameters and hardware.

**Fig 8.**
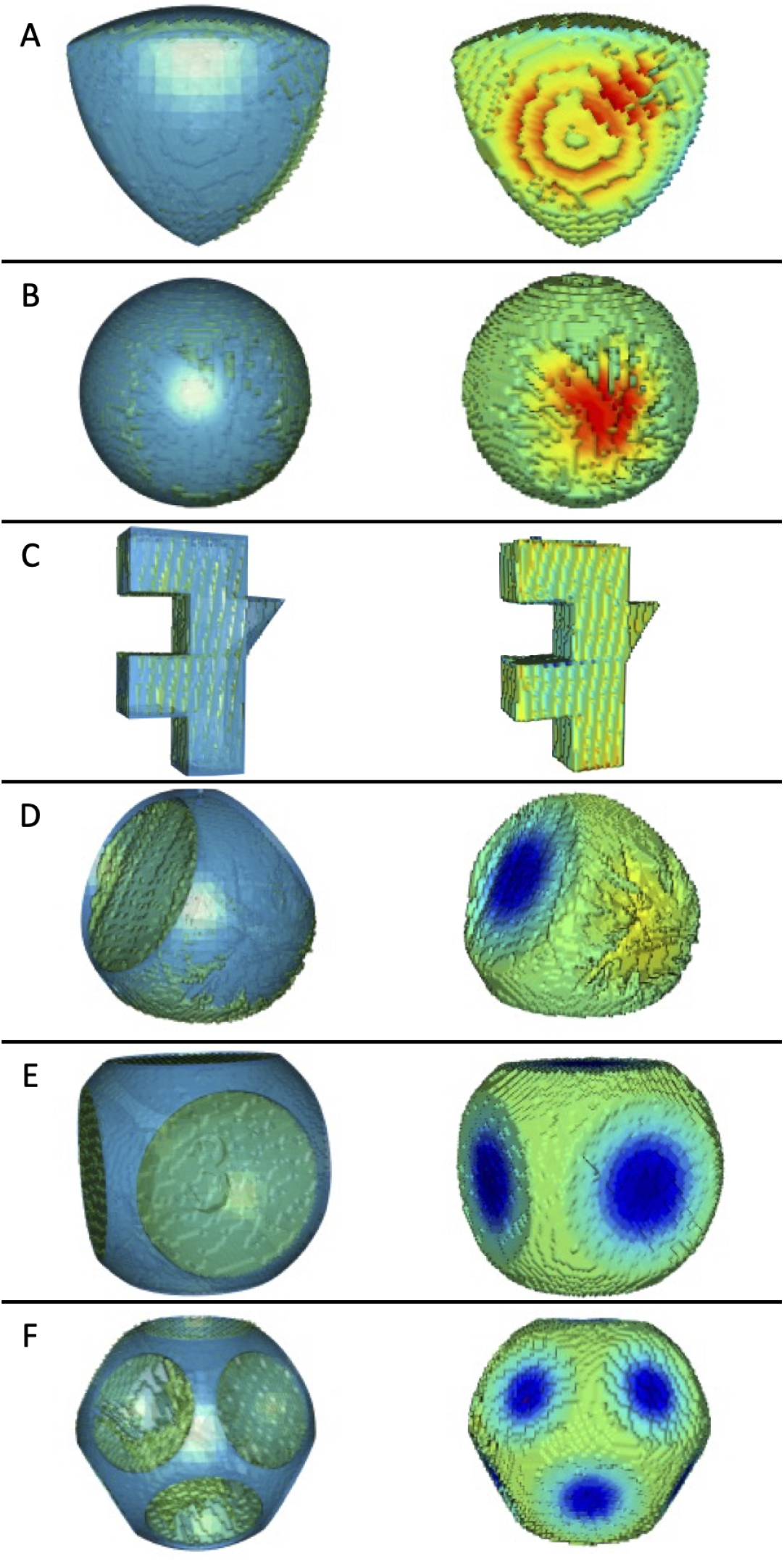
Reconstructions of six ground-truth objects. (Left Column) Reconstructed model (green) overlaid by ground-truth (blue) following ICP alignment. (Right column) Heat map showing difference between reconstructed model and ground-truth object. Red represents regions where the ground-truth is larger than the model. Blue represents regions where the ground-truth is smaller than the model. Teal represents regions where there is no difference between ground-truth and model. (A) A representative tetrahedron (Tetra_1), (A) a representative sphere (Sphere_1) (A) the smaller “F” shaped object, (D) the three sided die, (E) the six sided die, and (F) the twelve sided die. Only 1mm resolution models are shown. **Best viewed in color**.

**Table 1.** Accuracy metrics, including RMSE, difference in major axis length, and ratios of volume and surface area, from two experiments with eleven ground-truth objects. Differences in mm and ratios are reported between model and ground-truth object. Die_3, Die_6, and Die_12 have local concavities, while the other objects do not.

| Object | Res<sup>a</sup> | RMSE<sup>b</sup> | $\delta_X^c$ | $\delta_Y^d$ | $\delta_Z^e$ | $\delta_{Vol}^f$ | $\delta_{SA}^g$ |
| --- | --- | --- | --- | --- | --- | --- | --- |
| Tetra_2 | 1 | 0.62 | -0.12 | -0.14 | -0.14 | 0.97 | 1.20 |
|  | 2 | 0.76 | 0.04 | 1.17 | -1.17 | 0.96 | 1.18 |
| Tetra_3 | 1 | 1.17 | 0.61 | 0.21 | -0.93 | 0.96 | 1.24 |
|  | 2 | 1.21 | 1.32 | 0.35 | 0.97 | 0.95 | 1.23 |
| Tetra_4 | 1 | 1.62 | -0.29 | -0.83 | 0.05 | 0.96 | 1.26 |
|  | 2 | 1.69 | 1.51 | 0.31 | 0.06 | 0.95 | 1.25 |
| Sphere_2 | 1 | 0.97 | 0.47 | -0.26 | -0.53 | 0.96 | 1.23 |
|  | 2 | 1.07 | 0.97 | 0.21 | 1.38 | 0.95 | 1.21 |
| Sphere_3 | 1 | 1.54 | -3.27 | -1.01 | -1.25 | 0.96 | 1.27 |
|  | 2 | 1.57 | -4.74 | -1.07 | -1.20 | 0.95 | 1.26 |
| Sphere_4 | 1 | 2.61 | -0.70 | -1.70 | -1.81 | 0.95 | 1.33 |
|  | 2 | 2.70 | 0.33 | -1.19 | -0.90 | 0.94 | 1.33 |
| F_2 | 1 | 0.84 | 0.73 | -0.55 | 0.84 | 0.98 | 1.13 |
|  | 2 | 1.03 | 1.10 | 0.75 | 1.17 | 0.98 | 1.12 |
| F_3 | 1 | 0.66 | 4.66 | 0.82 | 1.93 | 1.01 | 1.16 |
|  | 2 | 0.79 | 4.11 | 1.45 | 2.09 | 1.00 | 1.15 |
| Die_3 | 1 | 7.05 | -3.16 | -9.64 | 5.68 | 1.17 | 1.13 |
|  | 2 | 6.64 | -2.70 | -5.18 | 1.14 | 1.16 | 1.10 |
| Die_6 | 1 | 6.31 | 8.71 | 2.45 | 6.52 | 1.29 | 1.03 |
|  | 2 | 6.00 | 9.77 | 8.25 | 7.97 | 1.28 | 1.03 |
| Die_12 | 1 | 9.34 | 0.28 | -4.21 | 8.61 | 1.20 | 1.02 |
|  | 2 | 8.70 | 0.27 | -2.02 | 9.57 | 1.20 | 1.00 |

<sup>a</sup>Reconstruction resolution in mm.
<sup>b</sup>RMSE of the model surface against the ground-truth surface mm.
<sup>c</sup>Difference between the X axis length of the model and the ground-truth mm.
<sup>d</sup>Difference between the Y axis length of the model and the ground-truth mm.
<sup>e</sup>Difference between the Z axis length of the model and the ground-truth mm.
<sup>f</sup>Ratio of the volume ( $Vol^{1/3}$ ) of the model over the volume of the ground-truth.
<sup>g</sup>Ratio of the surface area ( $SA^{1/2}$ ) of the model over the SA of the ground-truth.

**Fig 9.**
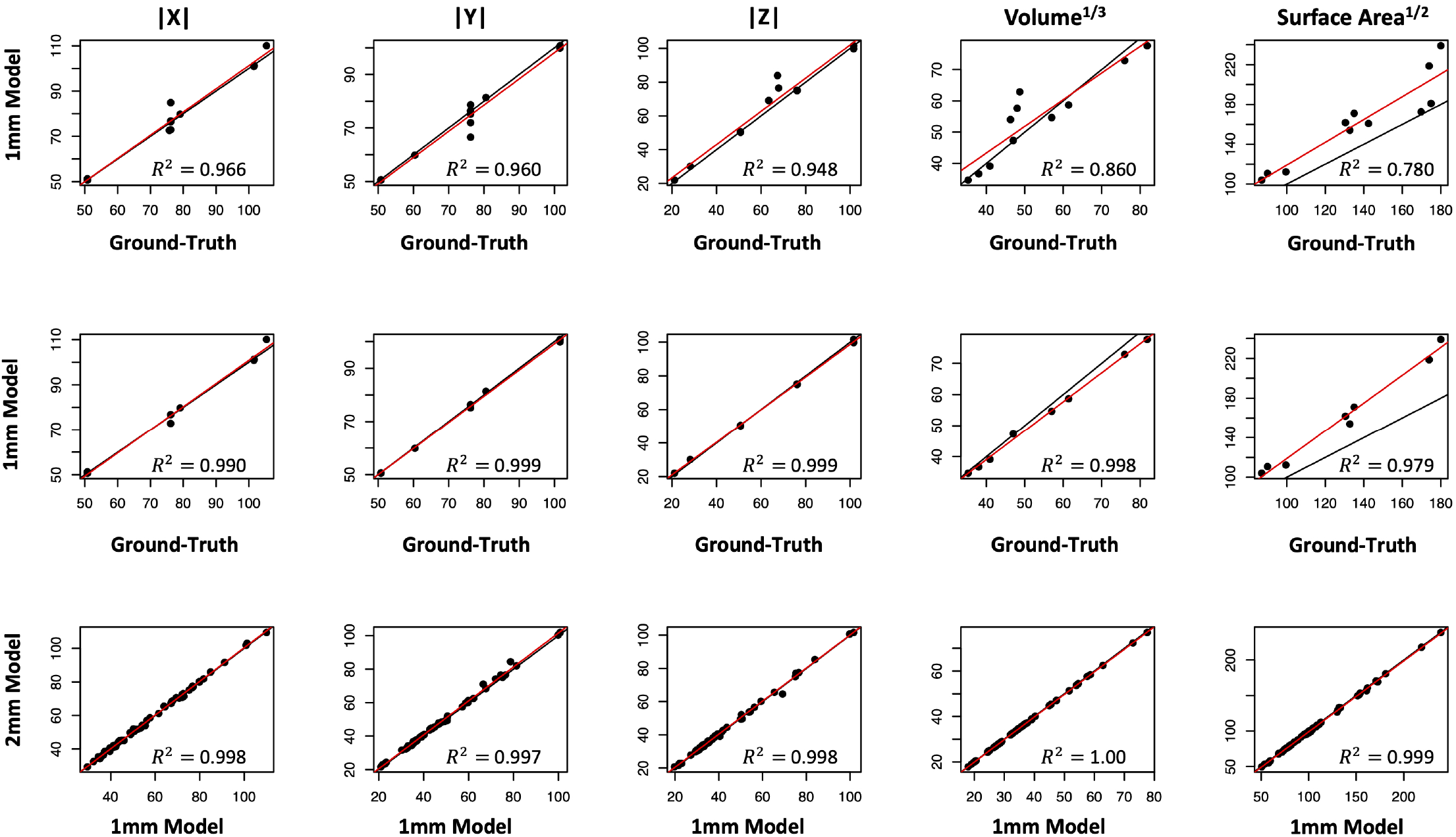
Ground-truth calibration experiment results. *In silico* measurement of reconstructed 3D models and ground-truth objects. Measurements include length of primary (X), secondary (Y), and tertiary (Z) axes, the cube root of the volume (Vol^1*/*3^), and the square root of the surface area (SA^1*/*2^). (Top row) All 1mm reconstructed models against ground-truth objects including the three dice, (Middle row) 1mm reconstructed models against ground-truth objects excluding the three dice, and (Bottom row) all reconstructed fruit models in 1mm (*x* axis) and 2mm (*y* axis). All measurements are reported in mm. The adjusted *R*^2^ from linear regression is reported in the plot. The solid black line is the identity line. The solid red line is the linear regression of *y* regressed onto *x* identity line.

The image calibration, segmentation, and reconstruction steps then proceed remotely following data organization and storage, which is an important consideration in practice. Using the 1 mm experiment to compute average run times, over all objects it took on average 27 seconds for calibration, 33.4 seconds for segmentation, and 413 seconds (6:53 minutes) for reconstruction, for an average total time of 7:53 minutes. All of these run times steps included load and write times of results and were generated on a workstation with one 12-core 2.70GHz processor and 192 GB of RAM.

The calibration, segmentation, and reconstruction steps are automated, and each of those steps are parallelized to some extent. Once the bounding box size was determined, the whole directory of samples was processed with a program that called each of the calibration, segmentation, and reconstruction steps, and was not supervised other than starting the process. Still, with new configurations or objects, or in case of failure, examining the output of each of the steps can indicate where there are problems, such as in the case of calibration or segmentation, on which the reconstruction step depends. For instance, in this hardware setup the calibrated camera poses should form a ring as in Figure 4. Accurate segmentations may be problematic for some objects, such as in Figure 8(F), the segmentation had false negatives at the bottom of the pear. Consequently, that part of the fruit is not reconstructed.

### Quantitative assessment of ground truth samples

In general, our approach performed very well on the groundtruth examples (Table 1) and only failed in ways that are were expected given the assumptions and constraints of our system^1^. Major deviations between reconstruction and groundtruth in the major axis are typically small (maximum 4.74 mm) and RMSE for the entire surface is *≤* 2.70 mm, for the models without concavities. We found a strong correlation *R*^2^ *≈* 0.99 for most measurements between the reconstructed models and the ground-truth objects without concavities (Figure 9). The surface area (SA) *R*^2^ was 0.979 for the models without concavities, and this *R*^2^ value is the lowest value for the traits we examined on objects without concavities. We found that SA of the reconstructed models were upwardly biased relative to the ground-truth objects (110-120%). This bias is most likely to do that fact that our models, which are made of voxels (Figure 5), have rough surfaces while the ground-truth objects are perfectly smooth (Figure 8). In general, the size measurement of these objects are very accurate, albeit imperfect, at both 1mm and 2mm resolutions (Figure 9).

We noticed minor segmentation false negative errors from shadows at the lower portion of the object in the images; in the reconstruction, these segmentation errors are realized as jagged portions where the printed object was attached to the pedestal, especially visible in Figure 8(A), (B), and (D). Reconstruction errors, resulting from small segmentation errors, do not have a large impact on the overall accuracy based on the metrics we assessed. However, large segmentation errors over multiple images will affect the reconstruction quality, such as in Figure 2F.

Our approach to reconstruction is unable to recover concavities, as demonstrated by the three spherical die examples (Figure 8D-F). The indexed faces of these models are sunken into the body of the model, resulting in multiple large depressions per die (Figure 7). As is clearly shown in Figure 8D-F, our reconstructions are more similar to a 3D convex hull, yielding a flat surface over the large concavities in the true models. This is reflected by the rows corresponding to the three die in Table 1. In these cases, the Volume is 115-130% greater than the ground-truth model. In general this is not an issue for types of fruit that do not have concavities.

### Qualitative assessment

We found that our platform and approach to reconstruction is both quantitatively accurate (Table 1; Figure 9), as well as visually accurate in most cases (Figures 2 and 8). For the peppers, grapes, strawberries, and potatoes, we found no systematic errors in reconstruction. However, the Anjou pears were troublesome to segment leading to the bottom half of the models being severely deformed. The reason for the segmentation error is likely the color variation of the pear and the use of a general segmentation approach that worked without extensive tuning for the whole set of samples. However, if a researcher were to have a large batch of objects with particular color features, fine tuning the user/session specific parameters for segmentation is important for yielding accurate models. Segmentation errors of this severe type appeared in 3 out of 59 objects that we imaged and the rest of the models appear to reflect the physical objects that were imaged.

## Discussion

We have described a low-cost ($1,600 USD), highthroughput (9 s data acquisition), modular reconstruction system that can be used in lab settings or in the field on a table, with a fast data acquisition speed of 9 seconds per object. We will discuss several design decisions that lead to flexibility. The use of consumer grade materials results in a relatively inexpensive system; multiple systems could be built and increase sample throughput during high-volume times of the year. This means that larger experiments can be executed enabling more robust studies. Our system is modular, allowing users with different interests to experiment and explore different cameras, sensors, or lights. This system is easily repaired and replaced if any damages are incurred by the hardware components.

The system has short session times and it only takes 9 seconds to acquire images on a single sample, regardless of the number of cameras. In fact, we found that were frequently rate limited by the write speed from the cameras on board cache to the SD card. More often than not, the next object was prepped around the same time that the cameras had cleared their on-board cache.

This system calibrates the camera from the image data acquired for the samples. The calibration is an absolute (as opposed to relative one, with an unknown scale factor), so the physical units of voxels are known.

## Key assumptions and considerations

We highlight some key assumptions of the methodologies used in our system that are important for those considering it for a range of objects not treated in this paper.

### Shape classes

Users who want to accurately represent locally concave shapes — shapes with egg-shaped depressions as demonstrated by the 3, 6, and 12-sided die in our calibration objects (Table 1; Figure 9; Figure 7)— will need to substitute some portions of this system to recover such features. Shape from Silhouette is not able to recover locally concave regions. However, most of the types of objects we envisioned imaging with this system, fruits and tubers, happen to be mostly non-concave.

### Large or fragile objects

The stepper motor is non-continuous and takes “steps” to provide rotation which causes vibrations through the object. When objects are unbalanced, those vibrations can cause movement of the object out of the center of the scene between different frames. Users should pay close attention to lateral movement of the object during rotation. Similarly, if the larger fruit are of interest, some additional modifications will be required to stabilize the pedestal during rotation. If these issues are a concern, it may be beneficial to construct a multi camera network that surrounds the target (Tabb et al., 2019) or a platform that enables the camera to move easily around a fixed target (Sandhu et al., 2019). The same is true for rice panicles or maize tassels, because they are not rigid body objects and the vibrations of the stepper motor are more likely to lead to changes in relative position of parts of the object between frames causing issues during reconstruction. These types of objects are better suited for systems where the cameras move relative to the object or in a multi-camera network (Sandhu et al., 2019; Tabb et al., 2019).

### Measurement of chArUco markers

Third, accuracy is intimately tied to the measurement of the arUco and chArUco markers and any inaccuracies in those measurements will lead to systematic biases in the measurement of the 3D reconstruction. For example, if chArUco markers are declared to be 10mm, when they are really 20mm, all of the models will 2x smaller than the real object than they measured. Users should print all calibration targets (aruco or chArUco) and ground-truth samples with a high quality 2D or 3D printer to ensure sharp corners and well-defined edges. Further, users are encouraged to verify the proportions of the printed calibration targets with high-precision calipers prior to calibration.

### Segmentation quality

Model quality is directly linked to segmentation quality (Figure 6A and 6B) as we have mentioned throughout this paper. If an object is only partially segmented, and this false negative error happens in multiple frames, part of the model may end up distorted or completely missing (Figure 2F). It is vital, as in any system, that users examine the quality of reconstructions prior to measurement and go back to calibration and segmentation outputs to identify the source of errors. In this work, we chose one set of segmentation parameters that performed reasonably well for all objects, but we recommend that users perform segmentation with parameters optimized for their research samples and imaging conditions.

## Conclusions

In conclusion, we presented a phenotyping system for capturing, calibrating, and reconstructing 3D models of small-to-moderately sized fruit and tubers. The low-cost and reliance on consumer-grade materials makes it obtainable to almost any program; short session times allows researchers to increase the number of samples per hour, and high accuracy means that the digital representations will yield absolute measurements on objects that do not degrade over time, yielding a viable option for research programs interested in pursuing 3D fruit phenotyping.

## Data Availability

The input data, including the images and configuration files, is available from Zenodo in (Feldmann and Tabb, 2021a), 10.5281/zenodo.5155765. All 3D model results, at 1 and 2 mm resolution, are at the same dataset source.

The ground truth objects were modified from objects at Thingiverse.com, and each had a different license. For this reason, there are four different datasets for the ground truth objects:

- Tetrahedra: 10.5281/zenodo.5153992 (Feldmann and Tabb, 2021e)
- Spheres: 10.5281/zenodo.5154029 (Feldmann and Tabb, 2021d),
- Sphere Dice: 10.5281/zenodo.5155690 (Feldmann and Tabb, 2021c).
- F-object: 10.5281/zenodo.5155743 (Feldmann and Tabb, 2021b)

Code for triggering cameras from Raspberri Pi GPIO have been deposited in the public Github repository (https://github.com/mjfeldmann/CameraTrigger).

## Acknowledgements

The authors thank Scott Wolford at USDA-ARS-AFRS for 3D printing the ground truth objects and Steven J. Knapp for supporting the application to and the Horticulture and Agronomy Graduate Group at UC Davis for awarding the Henry A. Jastro Research Scholarship.

Any mention of trade names or commercial products in this publication is solely for the purpose of providing scientific information and does not constitute recommendation or endorsement by the United States Department of Agriculture. USDA is an equal opportunity provider and employer.

## Author Contributions

Conceptualization: MJF, AT Data curation: MJF, AT Formal Analysis: MJF, AT Funding Acquisition: MJF, AT Investigation: MJF, AT Methodology: MJF, AT Project administration: MJF, AT Resources: MJF, AT Software: MJF, AT Supervision: MJF, AT Validation: MJF, AT Visualization: MJF, AT Writing – original draft preparation: MJF, AT Writing – review & editing: MJF, AT

